# TAL1-mediated epigenetic modifications down-regulate the expression of GIMAP family in non-small cell lung cancer

**DOI:** 10.1101/739870

**Authors:** Zhongxiang Tang, Lili Wang, Pei Dai, Pinglang Ruan, Dan Liu, Yurong Tan

## Abstract

**Backgroud:** The study was designed to explore the role of GIMAP family in non-small cell lung cancer (NSCLC) and its possible expression regulation mechanisms using existing biological databases including encyclopedia of DNA elements (ENCODE), gene expression omnibus (GEO), and the cancer genome atlas (TCGA).

**Methods:** Lung squamous cell carcinoma (LUSC) and lung adenocarcinoma (LUAD) were used to evaluate the expression of GIMAPs in TCGA. Five NSCLC datasets were then selected from GEO for validation. RNA-seq and Chip-seq data from ENCODE and GEO were used to observe epigenetic modifications on the chromosomes of GIAMPs in NSCLC. We constructed protein-protein interaction (PPI) network to reveal the main interacting proteins of GIMAPs. We then analyzed the correlation and regulatory mechanism between TAL1 and the expression of GIMAP family. We also used Kaplan-Meier for survival analysis.

**Results:** All 7 genes in the GIMAP family were down-regulated in NSCLC. H3K4me3 and H3K27ac disappeared, while H3K27me3 increased on the chromosome of this family. The expression of TAL1 was positively correlated with the expression of GIMAPs. sh-TAL1 significantly down-regulated the expression of GIMAP family through epigenetic modification. High expression of GIMAPs and TAL1 was found to be associated with a good prognosis of NLCSC.

**Conclusion:** The downregulation of TAL1 caused the disappearance of H3K27ac and H3K4me3. It also caused an increase in H3K27me3 on the GIMAPs gene, eventually leading to the overall downregulation of the GIMAP family genes.

## Backgroud

Lung cancer is a malignant disease of the lungs that has threatened human health for a long time now. It is stipulated that, for every four patients who die in the United States from cancer, one of them is a lung cancer patient [1]. Worldwide, approximately more than 1,600,000 people die of lung cancer yearly [2]. There are several types of lung cancer. However, the most important type of lung cancer based on histological classification is non-small cell lung cancer (NSCLC), accounting for about 85% of all cases if lung cancer. The pathological subtypes of NSCLC mainly include adenocarcinoma, squamous cell carcinoma and large cell carcinoma [3]. Several years of efforts and research has yielded some progress in the treatment of lung cancer. However, the main treatment of lung cancer is still limited to surgical chemotherapy, and the 5-year survival rate is still very low. Therefore, there is an urgent need to identify more sensitive and specific biomarkers for predicting tumor recurrence and prognosis to guide the treatment of NSCLC.

The human GTPase of the immunity-associated protein (GIMAP) family is composed of seven genes including GIMAP1, GIMAP2, GIMAP4, GIMAP5, GIMAP6, GIMAP7 and GIMAP8. In humans, the GIMAP subfamily genes are located in a cluster at 7q361[4], encoding an evolutionarily conserved GTP-binding protein that is preferentially expressed in immune cells. In experimental models and human pathology, specific members have been shown to be involved in the development of lymphocytes or in association with inflammation and autoimmune diseases [5]. Consistent with this, GIMAP1 is essential for the survival of naive and activated B cells *in vivo* [6]. GIMAP 4 participates in Th cell subset-driven immune balance through its role in T cell survival [7]. Deletion of GIMAP5 promotes the development of pathogenic CD4^+^T cells and the development of allergic airway disease [8]. Similarly, GIMAP is a potential modifier in autoimmune diabetes, asthma and allergies [9]. In the latest report, aberrant activation of the GIMAP enhancer by oncogenic transcription factors in T-cell induces acute lymphoblastic leukemia [10].

GIMAP is largely expressed in immune cells. It is also abundantly present in lung tissues, but the role of GIMAP in lung cancer has not been sufficiently reported. This study aimed to explore the expression of GIMAP in lung cancer using bioinformatics methods, and to determine its possible mechanism of low expression. It also seeks to add a key puzzle to the mechanism of NSCLC and to provide a new target for the diagnosis and treatment of NSCLC.

## Material and methods

### TCGA data visualization analysis

The cancer genome atlas (TCGA) database online analysis tool (http://gepia.cancer-pku.cn/) was used to analyze the expression of GIMAP family genes and T-cell acute leukemia 1(TAL1) by lung squamous cell carcinoma (LUSC) and lung adenocarcinoma (LUAD) in the NSCLC (LUAD tumor sample 483, normal sample 347; LUSC tumor sample 486, normal sample 338). Correlation function was then applied to analyze the correlation between TAL1 and GIMAP family gene expression.

### GEO microarry data

Five mRNA gene expression profiles GSE19804, GSE33532, GSE27262, GSE101929 and GSE21933 were downloaded from the National Center for Biotechnology Information Gene Expression Omnibus (GEO, https://www.ncbi.nlm.nih.gov/geo/) database. GSE101929 included 33 tumor samples and 33 paired non-tumor samples. GSE33532 consisted of 80 tumor samples and 20 corresponding normal samples. GSE27262 consisted of 25 tumor samples and 25 normal samples. GSE19084 consisted of 60 tumor samples and 60 normal samples while GSE21933 consisted of 21 tumor samples and 21 normal samples. Raw microarray data files (.CEL files) of the five datasets were downloaded from the GEO database. Background correction and quartile normalization were performed using the Robust Multichip Average (RMA) algorithm by the R package Affy [11].

Subsequently, the linear models for microarray data (limma) package in R were used to calculate the probability of probes being differentially expressed between cases and controls. The fold change (FC) and its logarithm value (log FC) were also determined. Corrected *p*-value < 0.05 and absolute ∣log2 fold change∣ > 1 were used to identify mRNAs that were differentially expressed significantly.

### CHIP-seq analysis

Chip-seq analysis of H3K27ac was performed using A549 (ENCODE: ENCFF256RBI), PC-9 (GSE96366), IMR-90 (ENCODE: ENCFF663RRL), normal lung (ENCFF105RLH) and left upper lobe (GSE101265). Chip-seq analysis of H3K4me3 was performed on A549 (GEO: GSE91218), PC-9 (GEO: GSE96170), IMR-90 (ENCODE: ENCFF721URL), normal lung (GEO: GSM1227065) and left upper lobe (GEO: GSE101033). A549 (GEO: GSE91306), PC-9 (GEO: GSE96468), IMR-90 (ENCODE: ENCFF806SLL) and normal lung tissue (GEO: GSM669969) were subjected to the Chip-seq analysis of H3K4me1. Chip-seq analysis of H3K27me3 was performed on A549 (GEO: GSM1003455), PC-9 (GEO: GSE96341), IMR-90 (ENCODE: ENCFF823FAY) and normal lung tissue (GEO: GSM706852).

### RNA-seq analysis and sh-TAL1

A549 (GEO: GSE93446), IMR-90 (GEO: GSE90257), left lung (GEO: GSM1101685), right lung (GEO: GSM1101692), lung (GEO: GSM1010946), left upper lobe (GEO: GSE88254) were used to perform RNA-seq. GSE72299 was used to perform sh-TAL1 analysis.

### PPI network construction

We used TRING database https://string-db.org/ with PPI output as constructed on the website. All items of GIMAP interaction were selected, and Cytoscape_v3.6.0 was used to calculate cytohubba. Network graph was then constructed using the GSE19804, GSE33532, GSE27262, GSE101929 and GSE21933 microarry data.

### Survival analysis of GIMAP family

Using the online survival analysis tool (http://kmplot.com/analysis/), we selected start KM plotter for lung cancer, as well as GIMAPs and TAL1 for overall survival analysis

## Results

### GIMAP family genes expression is downregulated in NSCLC

We explored the expression profile of GIMAP family genes in LUSC and LUAD using the TCGA dataset by online analysis tool GEPIA database (http://gepia.cancer-pku.cn/) [11]. The GEPIA box plots of GIMAP family gene expression levels showed that GIMAP1, GIMAP2, GIMAP4, GIMAP5, GIMAP6, GIMAP7 and GIMAP8 were low expressed in LUAD and LUSC except that the downgrade of GIMAP2 was not significant compared to the other six genes in LUAD (Fig.1). However, the adjacent gene ZNF775 and TMEM176B to GIMAP family genes were not found to be altered.

**Fig. 1.**
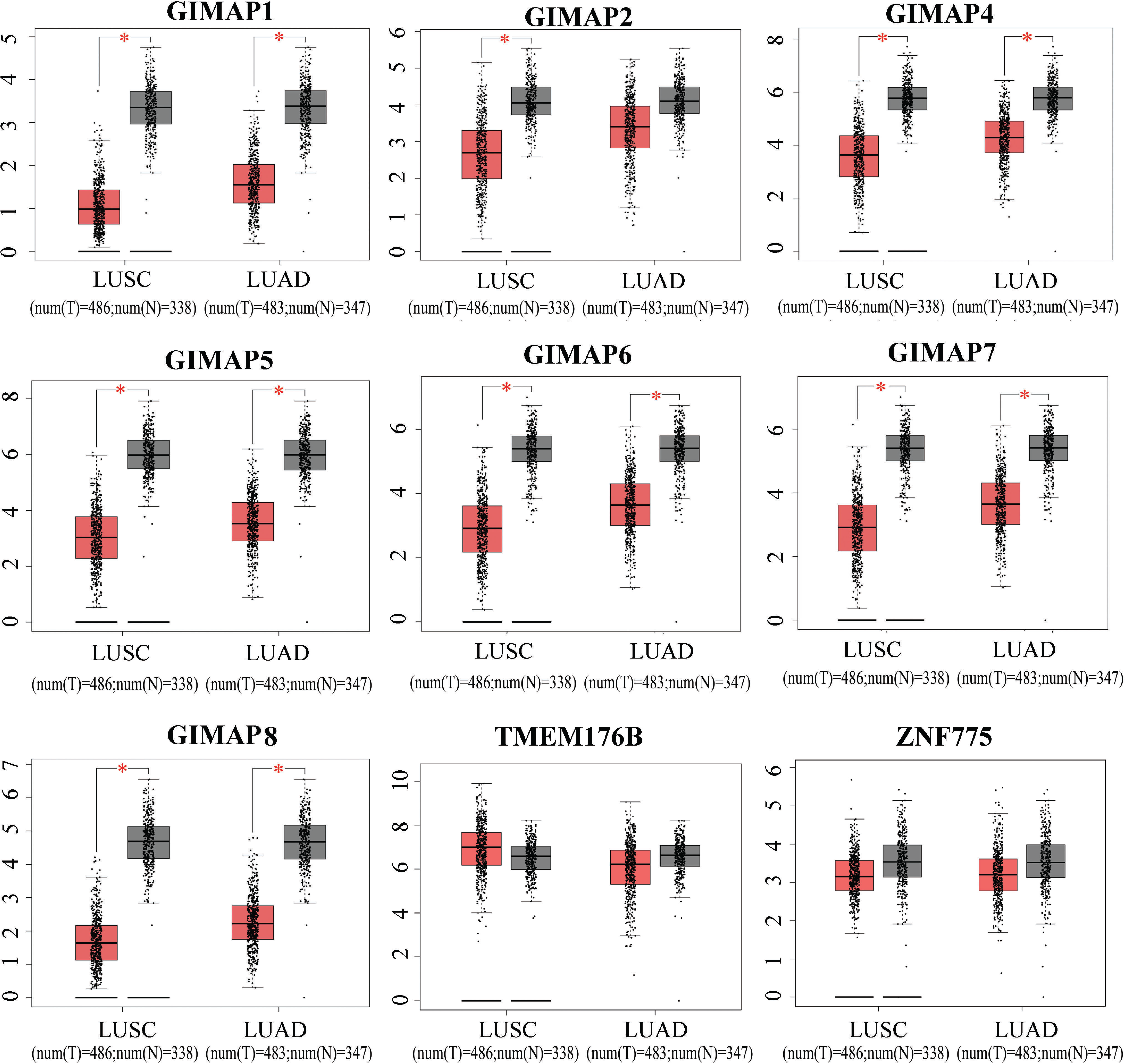
GEPIA boxed plots of GIMAP family in human LUSC, LUAD and normal lungs from TCGA. \**p* < 0.01 (LUSC: tumor=486, normal=338; LUAD: tumor=483, normal=347).

### GEO data confirms significant downregulation of GIMAPs in NLCSC

We set to confirm that GIMAPs are as consistently down-regulated as the TCGA database. Five NSCLC gene expression profiles (GSE19804, GSE33532, GSE27262, GSE101929, and GSE21933) were analyzed to confirm the GIMAP family genes that are expressed during tumorigenesis. The expression of mRNA in GIMAP family was significantly lower in NSCLC than in normal lung tissue. In addition, the amplitude of down-regulation of GIMAP2 was significantly weaker than that of the other GIMAP genes. The chromosome adjacent genes ZNF775 and TMEM176B did not exhibit any significant change (Fig. 2).

**Fig. 2.**
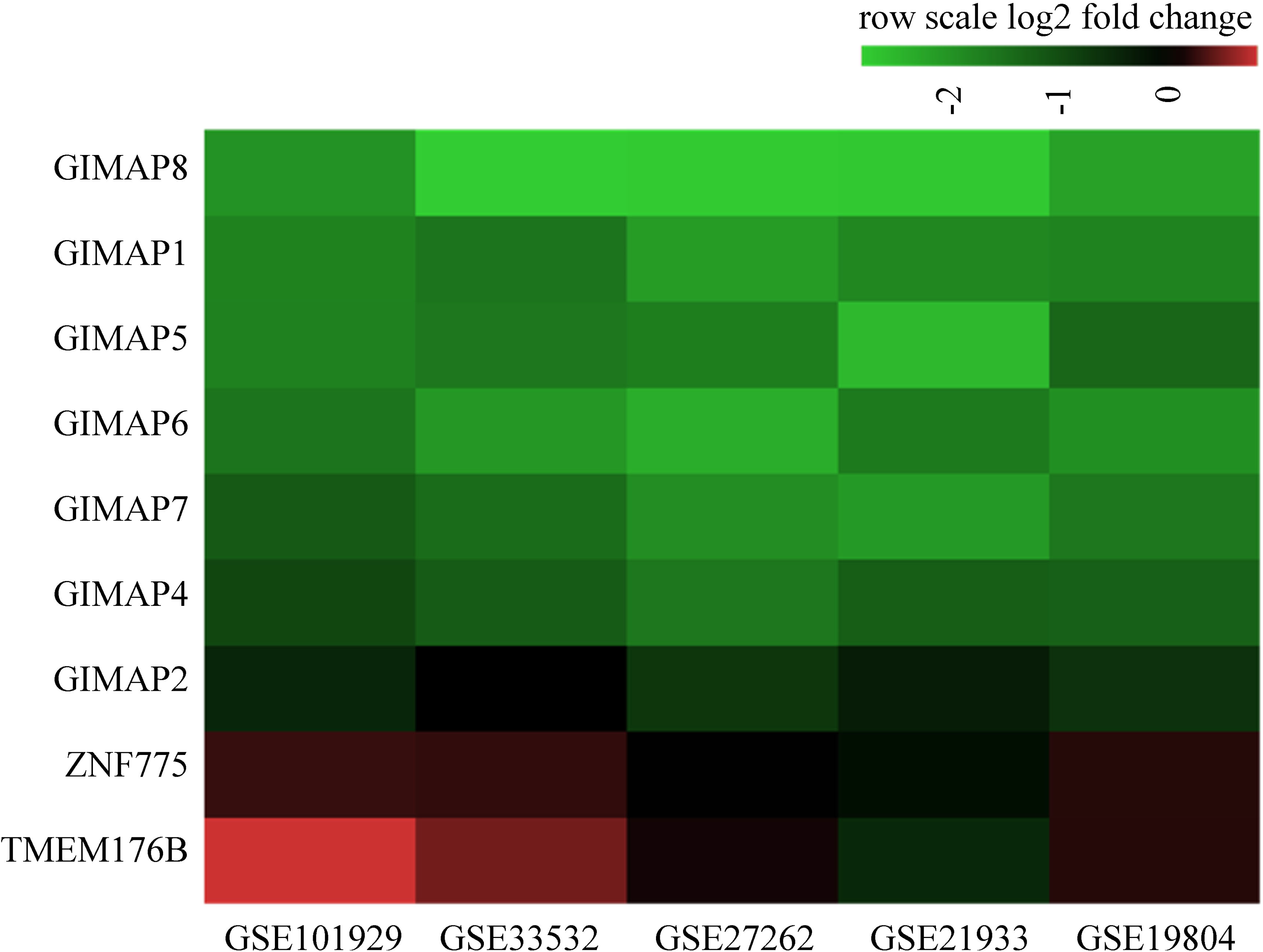
The GIMAP family gene was significantly down-regulated in five data sets, but there was no change in gene expression near the gene family.

### H3K4me3 and H3K27ac in GIMAPs disappeared and H3K27me3 enhanced in NLCSC

Epigenetic modification plays an important role in the transcriptional activation and participates in a number of life activities including differentiation, proliferation, apoptosis, immunity and tumor. We analyzed H3K4me1, H3K4me3, H3K27ac and H3K27me3 sites in GIMAPs in three cell lines A549, PC-9, IMR-90 of NSCLC, and made comparisons of RNA-seq at this position with normal lung tissue. The epigenetic modification of H3K27ac on the GIMAPs disappeared in A549, PC-9 and IMR-90, but it was apparent in the normal lung and the left upper lobe (Fig 3A).

**Fig. 3.**
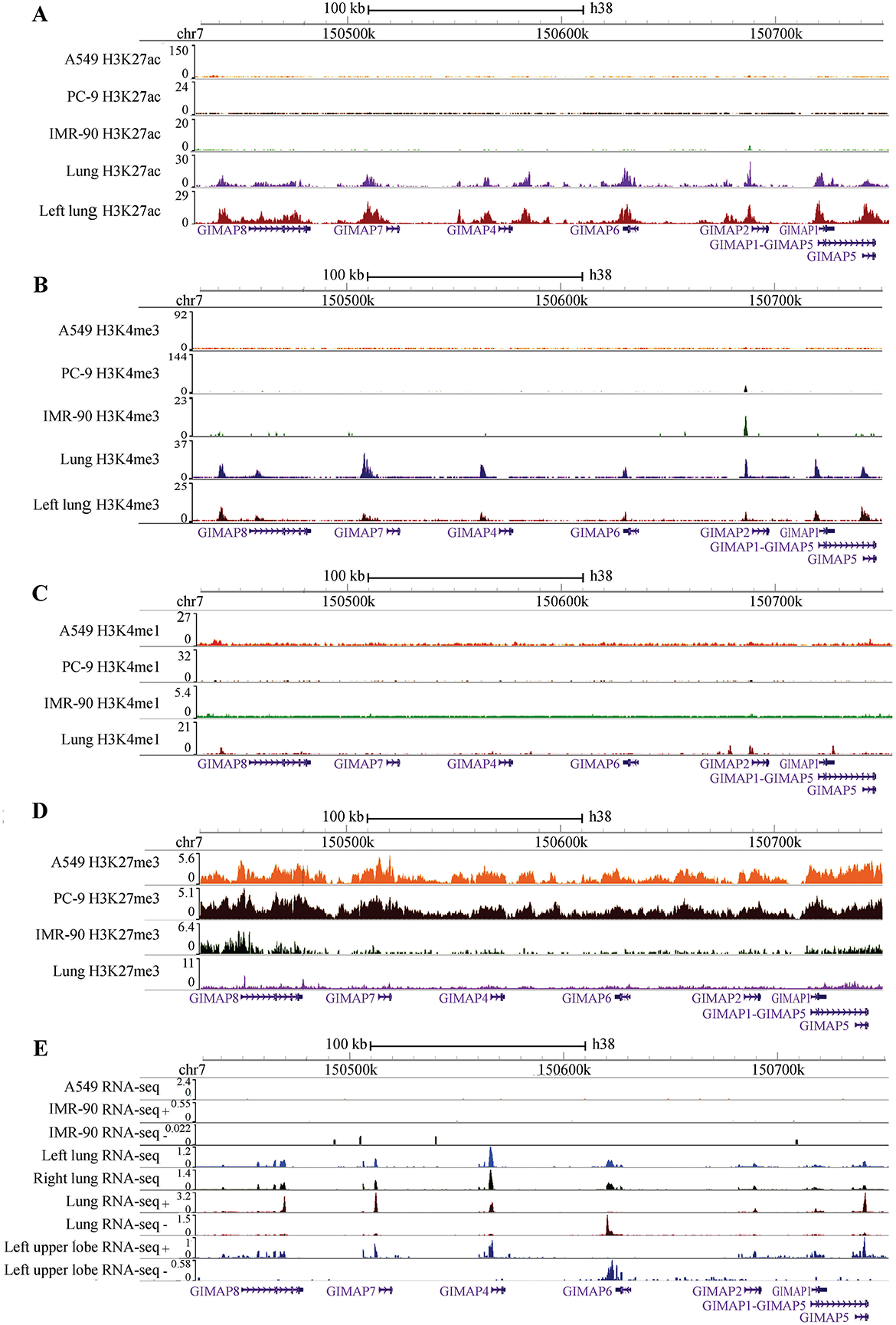
Epigenetic modifications of GIMAP family enhancer and promoter. A, H3K27ac; B, H3K4me3; C, H3k4me1; D, H3K27me3; E, RNA-seq of the GIMAPs.

However, the H3K27ac on ZNF775 and TMEM176B did not change. Moreover, the epigenetic modification of H3K4me3 at the GIMAPs disappeared in A549, PC-9 and IMR-90 too. However, there was a clear presence in the normal lung tissue and the left upper lobe (Fig 3B). However, the H3K4me3 on ZNF775 and TMEM176B did not change. It is interesting to note that H3K4me3 was still present at the GIMAP2 chromosomal location of A549, PC-9, and IMR-90. H3K4me1 on the GIMAPs exhibited no differences on A549, PC-9, IMR-90 and normal lung tissue (Fig 3C). However, in A549, PC-9 and IMR-90, H3K27me3 were significantly higher than that in normal lung tissue (Fig 3D). To confirm whether the expressions of GIMAPs in tumor cell lines were silenced, we performed RNA-seq of A549, IMR-90, left lung, right lung, lung and left upper lobe in TCGA and GEO. The data was consistent with the fact that the RNA-seq of the GIMAPs gene disappeared in A549 and IMR-90 but was apparent in normal tissues including left lung, right lung, lung and left upper lobe etc normal lung tissues (Fig 3E).

### TAL1 is the transcriptional regulator of GIMAPs in NLCSC

Five NSCLC gene expression programs (GSE19804, GSE33532, GSE27262, GSE101929, and GSE21933) were used for PPI analysis. We observed that GIMAP family can be interacted with FGR, CSF2RB, ABI3BP, ZMH, MNDA, VTA, VG204, T4, GD4, GZMA, P2RY13, and C1orf162 (Fig. 4A). TAL1 is a transcription factor involved in the transcription regulation of the GIMAP family and is related to epigenetic modification of genes. We further analyzed the relationship between the GIMAP family and TAL1 expression and observed that the expression of TAL1 is closely and positively related to seven GIMAP family members (Fig. 4B). Further, using shTAL1-seq, we found that, except GIMAP5, the expression of GIMAP family significantly decreased (Fig. 4C). Therefore, we speculated that in NLCSC, the expression of TAL1 is decreased, downregulating H3K27ac, H3K4me3 and promoting H3K27me3. This eventually leads to a down-regulation of the GIMAP family genes.

**Fig. 4.**
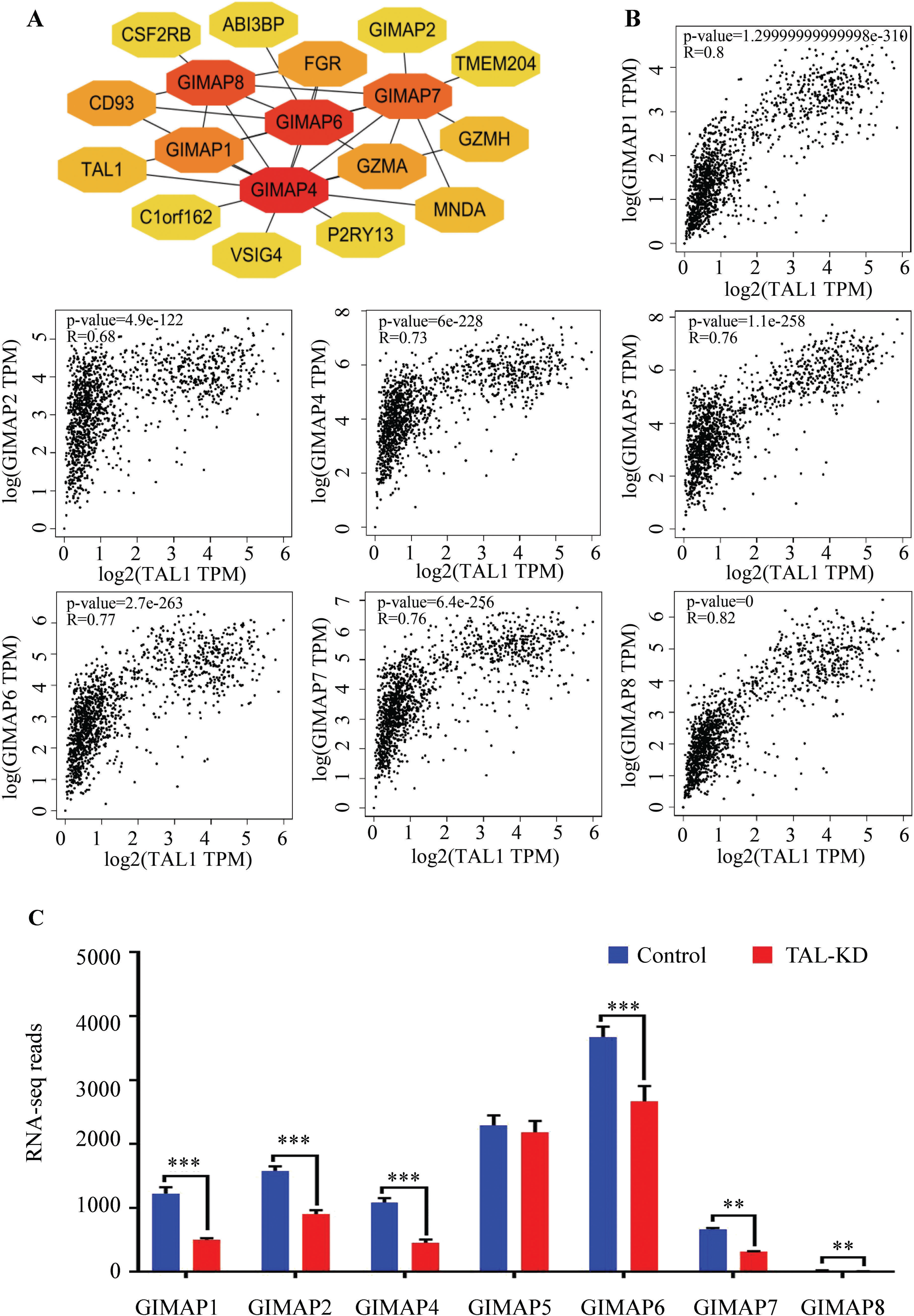
GIMAP family genes show significant positive correlations with the expression of transcription factor TAL1. A, PPI construction; B, the correlation between TAL1 and GIMAP family genes; C, shTAL1-seq found the expression of GIMAP family significantly decreased. *** *p*<0.001; ** *p*<0.01.

To further verify the regulatory mechanism of TAL1 on GIMAPs in NLCSC, Five NSCLC gene expression profiles (GSE19804, GSE33532, GSE27262, GSE101929, and GSE21933) were used for PPI analysis of TAL1. We found TAL1 may interact with PCAF, LSD1, HDAC1, HDAC2, TCF3 and EZH2. PCAF was found to be significantly increased, while LSD1, HDAC1, TCF3 and EZH2 were significantly up-regulated in lung cancer (F5A), which is consistent with the results in the TCGA database. Further correlation analysis of GIMAPs and PCAF, LSD1, HDAC1, HDAC2, TCF3 and EZH2 in the TCGA database showed that LSD1, HDAC1, TCF3 and EZH2 were significantly negatively correlated with GIMAPs, but PCAF were positively correlated. Therefore, we speculate that in NLCSC, TAL1 complex got involved in epigenetic modifications on H3K27ac, H3K4me3 and H3K27me3, eventually leading to a down-regulation of the GIMAPs gene.

**Fig. 5.**
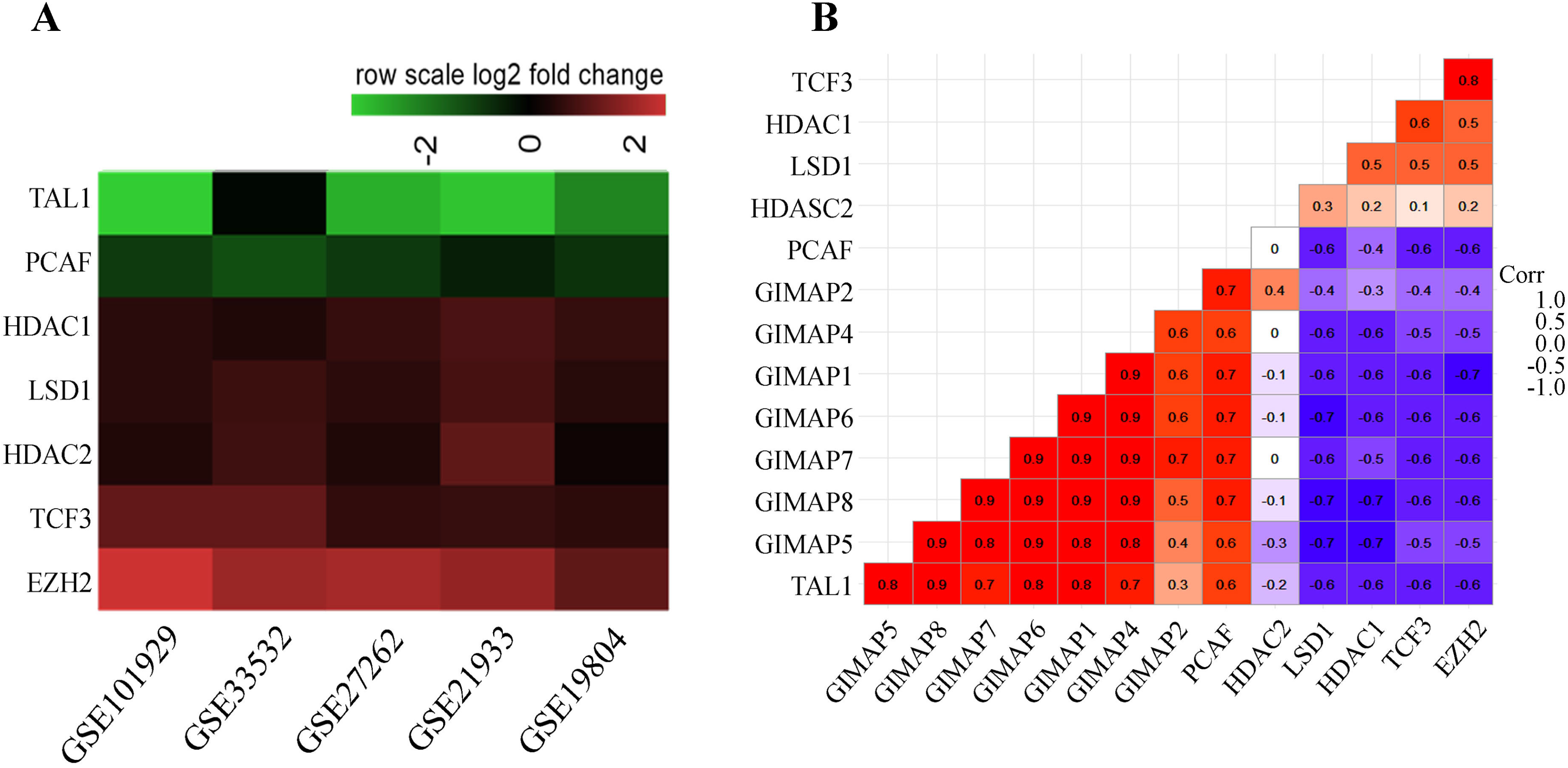
The regulatory mechanism of TAL1 on GIMAPs. A, the expression of TAL-1 interacted proteins in five datasets; B, the correlation between PCAF, LSD1, HDAC1, HDAC2, TCF3, EZH2 and GIMAP family genes.

### Low expression of GIMAP family genes is associated with bad prognosis

The survival analysis of GIMAP family and TAL1 was performed using the http://gepia.cancer-pku.cn/ online tool in LUAD. The results showed that high levels of *GIMAP1*, *GIMAP2*, *GIMAP4*, *GIMAP5*, *GIMAP*, *GIMAP7*, *GIMAP8* and *TAL1* mRNA expression were associated with better overall survival for LUAD patients (Fig.6).

**Fig. 6.**
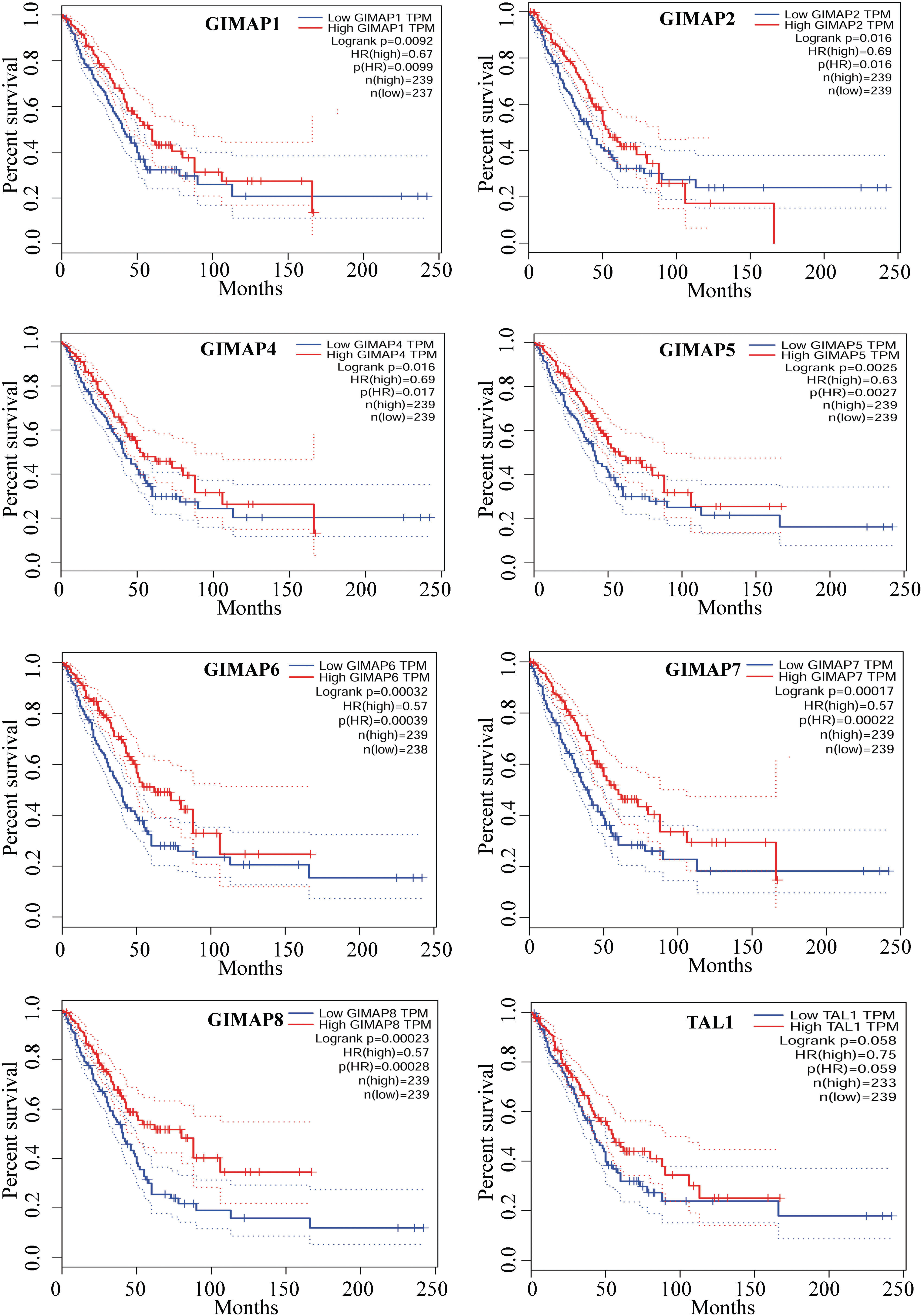
Overall survival analysis of GIMAP family and TAL1 in LUAD patients from TCGA database.

Finally, overall survival analysis of GIMAP family and TAL1was also performed using the Kaplan–Meier plotter online tool (http://kmplot.com/analysis/). The results also showed that high levels of all members of *GIMAP* family and *TAL1* mRNA was associated with better overall survival for lung cancer patients (Fig.7).

**Fig. 7.**
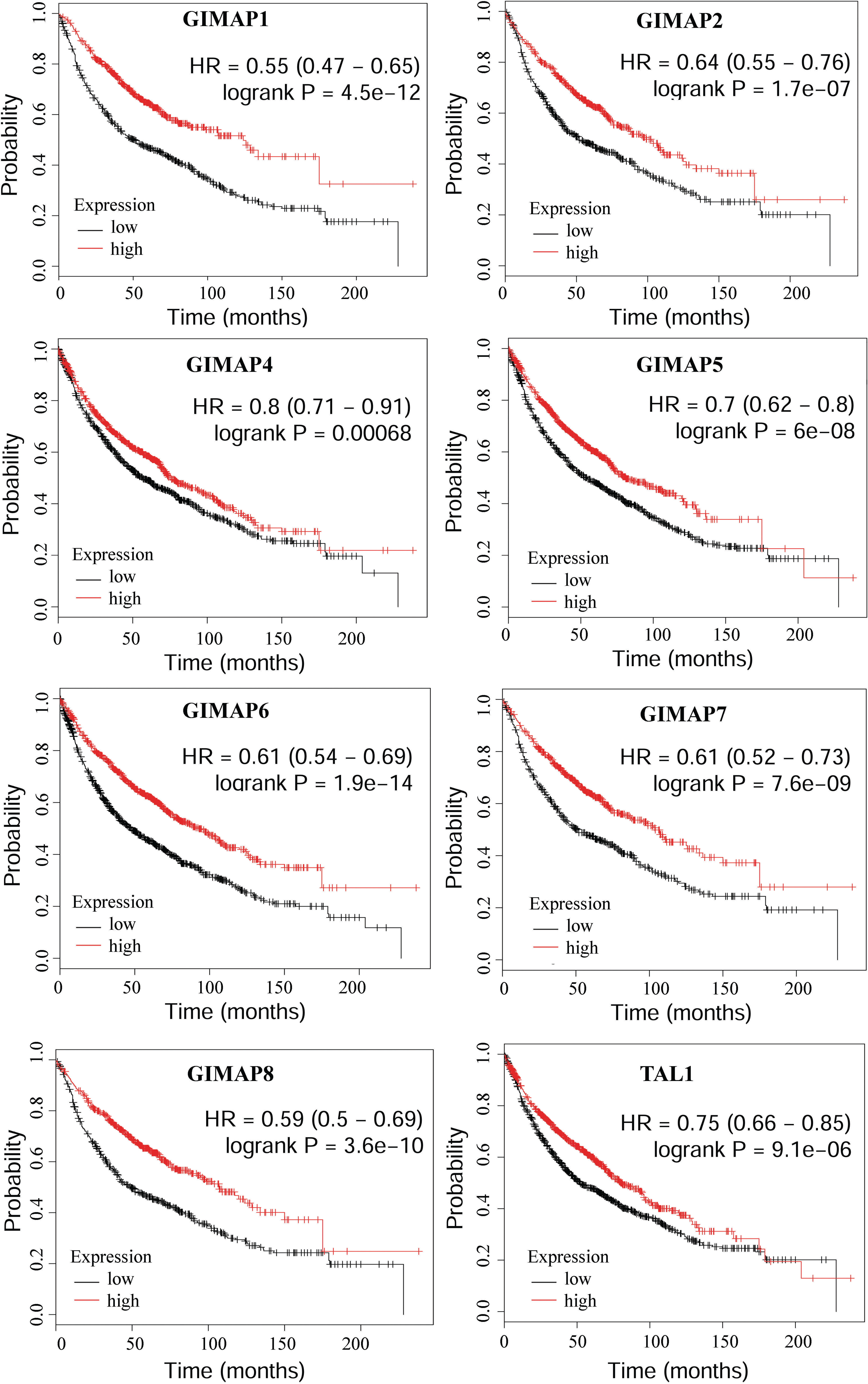
Prognostic value of GIMAP family and TAL1 in lung cancer patients from GEO database. Log-rank test was performed to evaluate the survival differences between the two curves.

## Discussion

Lung cancer is the leading cause of cancer death and NSCLC accounts for about 85% of all primary cancers of the lung. LUSC and LUAD are the two most common pathological types of NSCLC. Most patients with NSCLC present with an advanced state of the disease at the time of diagnosis. Platinum-containing chemotherapy can prolong the survival of patients, but the prognosis of patients with advanced NSCLC is still very poor, with a 5-year survival rate of less than 15%.

The *hGIMAPs* genes encoding these small GTPases belong to a family that is evolutionarily conserved in Arabidopsis, mice, rats and humans. These have been postulated to regulate apoptosis, particularly in the case of diseases such as cancer, diabetes and infection [7, 12]. In this study, we used the TCGA and GEO databases to report down-regulation of *hGIMAPs* in NLCSC and analyzed that low expression of all seven GIMAP genes was associated with poor prognosis, which is consistent with shiao et al [13]. GIMAPs are generally considered to be T cell receptor stress molecules and function primarily to exert anti-apoptotic effects in the immune system [14–16]. The current study found that GIMAPs are present in exosomes and significantly down-regulated in the PBMC cells of lung cancer patients [17–19]. Therefore, we surmised that down-regulation of GIMAPs in lung cancer led to deficiency of exogenous GIMAPs absorbed by T cells, leading to anti-tumor immune escape.

Epigenetic modifications include post-translational histones such as acetylation, phosphorylation, methylation, SUMOylation, and ADP-ribosylation and DNA methylation, which play a very complex regulatory role in transcription [20, 21]. These modifications can function independently or in concert, and rely on context-dependent ways to promote activation or inhibition of chromatin-mediated gene expression [22, 23]. Epigenetic modifications in GIMAP family genes were scanned in the present study, considering that the consistent downregulation of them may be related to gene epigenetic modification. H3K27ac and H3K4me3 are transcriptionally activated epigenetic modifications, usually expressed in the enhancer position of genes [24]. In addition to H3K4me3, H3K4me1 is also an epigenetic modification that activates transcription. It usually occurs at the promoter position of the gene, making the gene more accessible by transcription factors, thereby activating transcription [25]. H3K27me3 is generally considered to be a histone modification that inhibits transcriptional activation compared to H3K4me1, H3K4me3 and H3K27ac [26]. This present study indicated that the transcriptional activation of GIMAPs was mainly regulated by H3K4me3 and H3K27ac, and the transcriptional inhibition was regulated by H3K27me3. The GIMAP family was less affected by H3K4me1 in the lung.

Scientists have proposed the concept of "histone code", which consists of a specific pattern of histone modifications and can be decoded by chromatin-related proteins that recognize histone modifications. The histone code encoding and decoding process is in a state of dynamic equilibrium under normal conditions. Histone-modifying enzymes involved in encoding and decoding are often recruited to chromatin by DNA-binding transcription factors, ultimately leading to changes in chromatin structure and changes in gene expressio [27]. In order to further analyze the reasons for the down-regulation of GIMAP family genes, we used differentially expressed gene (DEG) to analyze PPI. The results showed that TAL1 was the upstream regulatory gene of GIMAP family, and is involved in epigenetic modifications. GIMAP and TAL1 expression were significantly positively correlated and shRNA-TAL1 results in down-regulating GIMAP expression. TAL1 is a member of the basic helix-loop-helix (bHLH) transcription factor family. Several studies have shown that TAL1 can act as a transcriptional activator or as a repressor in a context-dependent manner. The TAL1 transcription factor contains a threonine (Thr) residue and a serine (Ser) residue, which can be phosphorylated by protein kinase A (PKA), which can specifically disrupt the interaction between TAL1 and H3K4 demethylase LSD1, leading to increased H3K4 methylation and activation of target genes that have been suppressed in malignant hematopoiesis [28]. Moreover, in the process of erythroid differentiation, the interaction of TAL1 with the histone demethylase decreases, and the binding to the histone methyltransferase LSD1 increases, accompanied by H3K4 methylation [28]. Therefore, it is speculated that TAL1 may be a core factor in histone coding. During DNA transcription, TAL1 dynamically recruits histone acetyltransferases (P300/CBP-associated factor, PCAF), histone deacetylases (HDACs), histone methyl groups under normal conditions. In this study, PCAF was significantly down-regulated, and HDAC1 were significantly increased leading to decreased H3K27ac in lung cancer. At the same time, the histone demethylases EZH2 and LSD1 were significantly upregulated in this study, leading to decreased H3K4 methylation. The inhibitors of HDAC have shown the effect of inducing lung cancer cell death in clinical trial [29]. Histone methyltransferase inhibitors azacytidine and etinostat inhibit the expression of EZH2, thereby inhibiting NSCLC [30].

In conclusion, our results show that the expression levels of the GIMAP gene family are consistently down-regulated followed by the disappearance of H3K27ac and H3K4me3 and the increase of H3K27me3 on the GIMAPs in NSCLC. TAL1 was positively correlated with GIMAPs and shRNA-TAL1 led to the overall downregulation of the GIMAP family genes. Low expression of all seven GIMAP genes is associated with poor prognosis.

## Abbreviations

GIMAP: GTPase, IMAP family member
ZNF: zinc finger protein 775
TAL1: TAL bHLH transcription factor 1, erythroid differentiation factor
NSCLC: nonsmall-cell lung cancer
TMEM176B: transmembrane protein 176B
TCGA: The Cancer Genome Atlas
RAM: robust multichip average
GEO: Gene Expression Omnibus
ENCODE: Encyclopedia of DNA Elements
LUAD: Lung Adenocarcinoma
LUSC: Lung Squamous Cell Carcinoma
EZH2: enhancer of zeste 2 polycomb repressive complex 2 subunit
HDAC: Histone deacetylase
PCAF: lysine acetyltransferase 2B
LSD1: lysine demethylase 1A
TCF3: transcription factor 3
H3K4me3: trimethylation of lysine 4 on histone H3 protein subunit
H3K27ac: acetylation of lysine 4 on histone H3 protein subunit
H3K27me3: trimethylation of lysine 27 on histone H3 protein subunit

## Declarations

### Authors’ contributions

YT and LW designed this work. ZT and YT wrote the paper. ZT analyzed the data. PD, DL, and PR joined discussions. All authors have approved the present version of the manuscript and have agreed to be accountable for all aspects of the work regarding questions related to the accuracy or integrity of any part of the work. All authors read and approved the final manuscript.

#### Acknowledgements

Not applicable.

### Availability of data and materials

The datasets used and/or analyzed during the current study are available from the corresponding author on reasonable request.

### Consent for publication

Not applicable.

### Ethics approval and consent to participate

Not applicable.

### Publisher’s Note

Springer Nature remains neutral with regard to jurisdictional claims in published maps and institutional affiliations.

## References

1. Siegel RL, Miller KD, Jemal A: Cancer Cancer statistics, 2018. CA Cancer J Clin 2018, 68(1):7–30.

2. Herbst RS, Morgensztern D, Boshoff C: The biology and management of non-small cell lung cancer. Nature 2018, 553(7689):446–454.

3. Gerber DE, Minna JD: ALK inhibition for non-small cell lung cancer: from discovery to therapy in record time. Cancer Cell 2010, 18(6):548–551.

4. Filen S, Lahesmaa R: GIMAP Proteins in T-Lymphocytes. J Signal Transduct 2010, 2010:268589.

5. Ciucci T, Bosselut R: Gimap and T cells: a matter of life or death. Eur J Immunol 2014, 44(2):348–351.

6. Webb LM, Datta P, Bell SE, Kitamura D, Turner M, Butcher GW: GIMAP1 Is Essential for the Survival of Naive and Activated B Cells In Vivo. Journal of immunology (Baltimore, Md: 1950) 2016, 196(1):207–216.

7. Heinonen MT, Laine AP, Soderhall C, Gruzieva O, Rautio S, Melen E, Pershagen G, Lahdesmaki HJ, Knip M, Ilonen J et al. GIMAP GTPase family genes: potential modifiers in autoimmune diabetes, asthma, and allergy. Journal of immunology (Baltimore, Md: 1950) 2015, 194(12):5885–5894.

8. Patterson AR, Bolcas P, Lampe K, Cantrell R, Ruff B, Lewkowich I, Hogan SP, Janssen EM, Bleesing J, Khurana Hershey GK et al.: Loss of GTPase of immunity-associated protein 5 (Gimap5) promotes pathogenic CD4(+) T-cell development and allergic airway disease. J Allergy Clin Immunol 2019, 143(1):245–257 e246.

9. Heinonen MT, Kanduri K, Lahdesmaki HJ, Lahesmaa R, Henttinen TA: Tubulin-and actin-associating GIMAP4 is required for IFN-gamma secretion during Th cell differentiation. Immunol Cell Biol 2015, 93(2):158–166.

10. Liau WS, Tan SH, Ngoc PCT, Wang CQ, Tergaonkar V, Feng H, Gong Z, Osato M, Look AT, Sanda T: Aberrant activation of the GIMAP enhancer by oncogenic transcription factors in T-cell acute lymphoblastic leukemia. Leukemia 2017, 31(8):1798–1807.

11. Tang Z, Li C, Kang B, Gao G, Li C, Zhang Z: GEPIA: a web server for cancer and normal gene expression profiling and interactive analyses. Nucleic Acids Res 2017, 45(W1):W98–W102.

12. Kupfer R, Lang J, Williams-Skipp C, Nelson M, Bellgrau D, Scheinman RI: Loss of a gimap/ian gene leads to activation of NF-kappaB through a MAPK-dependent pathway. Mol Immunol 2007, 44(4):479–487.

13. Shiao YM, Chang YH, Liu YM, Li JC, Su JS, Liu KJ, Liu YF, Lin MW, Tsai SF: Dysregulation of GIMAP genes in non-small cell lung cancer. Lung Cancer 2008, 62(3):287–294.

14. Nitta T, Nasreen M, Seike T, Goji A, Ohigashi I, Miyazaki T, Ohta T, Kanno M, Takahama Y: IAN family critically regulates survival and development of T lymphocytes. PLoS Biol 2006, 4(4):e103.

15. Nitta T, Takahama Y: The lymphocyte guard-IANs: regulation of lymphocyte survival by IAN/GIMAP family proteins. Trends Immunol 2007, 28(2):58–65.

16. Datta P, Webb LM, Avdo I, Pascall J, Butcher GW: Survival of mature T cells in the periphery is intrinsically dependent on GIMAP1 in mice. Eur J Immunol 2017, 47(1):84–93.

17. Kowal J, Arras G, Colombo M, Jouve M, Morath JP, Primdal-Bengtson B, Dingli F, Loew D, Tkach M, Thery C: Proteomic comparison defines novel markers to characterize heterogeneous populations of extracellular vesicle subtypes. Proceedings of the National Academy of Sciences of the United States of America 2016, 113(8):E968–977.

18. Skogberg G, Gudmundsdottir J, Van der Post S, Sandstrom K, Bruhn S, Benson M, Mincheva-Nilsson L, Baranov V, Telemo E, Ekwall O: Characterization of human thymic exosomes. PloS one 2013, 8(7):e67554.

19. Perez-Hernandez D, Gutierrez-Vazquez C, Jorge I, Lopez-Martin S, Ursa A, Sanchez-Madrid F, Vazquez J, Yanez-Mo M: The intracellular interactome of tetraspanin-enriched microdomains reveals their function as sorting machineries toward exosomes. The Journal of biological chemistry 2013, 288(17):11649–11661.

20. Kouzarides T: SnapShot: Histone-modifying enzymes. Cell 2007, 131(4):822.

21. Rando OJ, Chang HY: Genome-wide views of chromatin structure. Annu Rev Biochem 2009, 78:245–271.

22. Darilmaz Yuce G, Ortac Ersoy E: [Lung cancer and epigenetic modifications]. Tuberk Toraks 2016, 64(2):163–170.

23. Berger SL: Histone modifications in transcriptional regulation. Curr Opin Genet Dev 2002, 12(2):142–148.

24. Dahl JA, Jung I, Aanes H, Greggains GD, Manaf A, Lerdrup M, Li G, Kuan S, Li B, Lee AY et al. Broad histone H3K4me3 domains in mouse oocytes modulate maternal-to-zygotic transition. Nature 2016, 537(7621):548–552.

25. Yan J, Chen SA, Local A, Liu T, Qiu Y, Dorighi KM, Preissl S, Rivera CM, Wang C, Ye Z et al.: Histone H3 lysine 4 monomethylation modulates long-range chromatin interactions at enhancers. Cell Res 2018, 28(2):204–220.

26. Agger K, Cloos PA, Christensen J, Pasini D, Rose S, Rappsilber J, Issaeva I, Canaani E, Salcini AE, Helin K: UTX and JMJD3 are histone H3K27 demethylases involved in HOX gene regulation and development. Nature 2007, 449(7163):731–734.

27. Consortium EP: An integrated encyclopedia of DNA elements in the human genome. Nature 2012, 489(7414):57–74.

28. Li Y, Deng C, Hu X, Patel B, Fu X, Qiu Y, Brand M, Zhao K, Huang S: Dynamic interaction between TAL1 oncoprotein and LSD1 regulates TAL1 function in hematopoiesis and leukemogenesis. Oncogene 2012, 31(48):5007–5018.

29. Brazelle W, Kreahling JM, Gemmer J, Ma Y, Cress WD, Haura E, Altiok S: Histone deacetylase inhibitors downregulate checkpoint kinase 1 expression to induce cell death in non-small cell lung cancer cells. PloS one 2010, 5(12):e14335.

30. Komashko VM, Farnham PJ: 5-azacytidine treatment reorganizes genomic histone modification patterns. Epigenetics 2010, 5(3):229–240.

